# Using single-cell entropy to describe the dynamics of reprogramming and differentiation of induced pluripotent stem cells

**DOI:** 10.1101/2020.04.13.040311

**Authors:** Yusong Ye, Zhuoqin Yang, Meixia Zhu, Jinzhi Lei

## Abstract

Induced pluripotent stem cells (iPSCs) provide a great model to study the process of stem cell reprogramming and differentiation. Single-cell RNA sequencing (scRNA-seq) enables us to investigate the reprogramming process at single-cell level. Here, we introduce single-cell entropy (scEntropy) as a macroscopic variable to quantify the cellular transcriptome from scRNA-seq data during reprogramming and differentiation of iPSCs. scEntropy measures the relative order parameter of genomic transcriptions at single cell level during the process of cell fate changes, which show increase tendency during differentiation, and decrease upon reprogramming. Hence, scEntropy provides an intrinsic measurement of the cell state, and can be served as a pseudo-time of the stem cell differentiation process. Moreover, based on the evolutionary dynamics of scEntropy, we construct a phenomenological Fokker-Planck equation model and the corresponding stochastic differential equation for the process of cell state transitions during pluripotent stem cell differentiation. These equations provide further insights to infer the processes of cell fates changes and stem cell differentiation. This study is the first to introduce the novel concept of scEntropy to quantify the biological process of iPSC, and suggests that the scEntropy can provide a suitable macroscopic variable for single cells to describe cell fate transition during differentiation and reprogramming of stem cells.

## 1 Introduction

Induced pluripotent stem cells (iPSCs) are derived from skin or blood cells that have been reprogrammed back into an embryonic-like pluripotent state that enables the development of other types of differentiated cells. Reprogramming of iPSC is often induced through the induction of genes important for maintaining the essential properties of embryonic stem cells (ESCs), and the genomic transcriptions change during the processes of reprogramming and further differentiation. However, these processes are rather stochastic, and the molecular processes of cell fate changes remain unclear[10]. Recently, single-cell RNA sequencing (scRNA-seq) methods have allowed for the investigation of cellular transcriptome at the level of individual cells[1, 3, 11]. The technique of scRNA-seq enables us to better study the dynamics of reprogramming and differentiation of iPSCs, which can provide insightful information on the process of cell reprogramming[2, 7, 13]. Based on the microscopic state of genomic transcriptions provided by scRNA-seq data, we are able to identify the changes of marker genes expressions during the reprogramming process. Alternatively, a well defined macroscopic variable for the state of a cell is important for further understanding of the dynamic process.

Recently, a novel concept of single-cell entropy (scEntropy) was proposed to measure the order of cellular transcriptome profile from scRNA-seq data[6]. The scEntropy of a cell is defined as the information entropy of the difference in transcriptions between the cell and a predefined reference cell, and provides a straightforward and parameter-free macroscopic variable that can be used to quantify the process of early embryo development[6]. Here, we investigate whether scEntropy can be used to describe the reprogramming and the differentiation processes of iPSCs. We introduce the concept of scEntropy to quantify the states of individual cells along with the reprogramming and differentiation processes. The scEntropy shows obvious increase trends during differentiation, and decrease tendencies during reprogramming. Thus, scEntropy can be served as a pseudo-time of the process, according to which we identify the genes that show expressions correlated with the scEntropy, the genes form potent markers of cell fate changes. We also construct phenomenological Fokker-Planck equation model and the associated stochastic differential equations for the plasticity process of stem cells.

## 2 Results

### 2.1 scEntropy to describe the process of differentiation and reprogramming

The scEntropy was proposed to measure the order of cellular transcription from scRNA-seq data with respect to a reference level; larger entropy means lower order in the transcriptions[6]. Given an *N × M* gene expression matrix with *N* cells and *M* genes, and the gene expression vector **r** of the reference cell. Let **x_*i*_** (*i* = 1,…,*N*) the gene expression vector of the *i*^th^ cell. Calculation of the scEntropy of **x_*i*_** with reference to **r**, *S*(**x_*i*_|r**), includes two steps[6]: (1) calculate the difference between **x_*i*_** and **r**

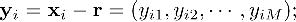

(2) the entropy ***S***(**x_*i*_|r**) is given by the information entropy of the signal sequence **y_*i*_**, *i.e*.,

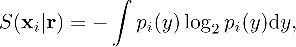

where *p_i_*(*y*) is the distribution density of the components *y_ij_* in **y_*i*_**. From the definition, the reference cell **r** is a variable in defining the entropy *S*(**x_*i*_|r**), which means the baseline transcriptome with the minimum entropy of zero.

To study the dynamics of reprogramming and differentiation of iPSCs, we can refer the state of iPSC as the reference cell in defining the scEntropy, the resulting scEntropy gives the relative information entropy of each cell with respect to the state of iPSC.

To illustrate the application of scEntropy, we apply scEntropy to investigate the kidney organoid differentiation from human pluripotent stem cells[12]. Time-series scRNA-seq data of totally 9,190 cells were obtained from separate time points (days 0, 7, 12, 19, and 26) during the differentiation process (GSE118184). To calculate the scEntropy, we take the average of gene expressions of cells at day 0, so that scEntropy gives the relative transcription order with respect to the state of human pluripotent stem cells. Here, the scEntropies are calculated from top 500 high variance genes using Find Variable Features in Seurat.

The scEntropies of the sequenced cells at each day are shown in Fig. 1A. We note that the cells are heterogeneous at each day so that the scEntropies of cells at each day distributed over a wide range from 4.8 to 5.8. And there is a tendency of increasing scEntropies along the differentiation process from day 0 to day 26. We further calculate the average and variance of the scEntropies at each day (Fig. 1B-C). The average scEntropy obviously increases during differentiation, and the variance maintains the same level during the differentiation process except for a low level variation on day 26.

**Figure 1:**
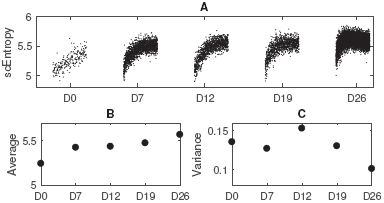
scEntropy dynamics during kidney organoid differentiation. **A**. scEntropies of the transcription of 9,190 cells from separate time points during kidney organoid differentiation from human pluripotent stem cells(GSE118184). The dots are scattered in each day for avoiding mixing and overlapping, and are ordered the same as the original data set. **B**. Time course of the average scEntropies of cells sequenced at each day. **C**. Variance of the scEntropies of cells sequenced at each day.

Next, we analyze the scEntropy of mouse cells during the reprogramming process from MEF to iPSC[4]. A total of 912 mouse cells were sequenced from 9 time points from to iPSC (GSE103221). We take the average gene expression vectors of all iPSCs as the reference cell, and calculated the scEntropies of all cells. The scEntropy is nearly unchanged in 8 days after induction, and obviously decreases upon further reprogrammed into iPSC (Fig. 2).

**Figure 2:**
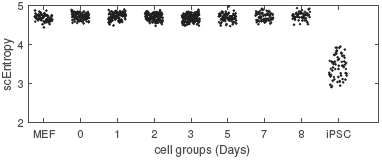
scEntropy of cells from 9 time points during the process of reprogramming from MEF to iPSC. The dots are scattered randomly in each day for avoiding mixing and overlapping.

While the concept of scEntropy was original introduced based on scRNA-seq data, it is also valid for bulk data sets. When **x_*i*_** represents the average transcription of a group of cells, the entropy *S*(**x_*i*_|r**) gives the relative transcriptional order of the average transcriptome of these cells with respect to the reference cell.

Here, we apply scEntropy to investigate the differentiation of iPSCs to cardiomyocytes based on a bulk data set of time course RNA-seq data. The data set includes 297 RNA samples, each from separate time points in 19 human Yoruba Hapmap cell lines (GSE122380)[9]. In experiments, the cells in one well were harvested and frozen every 24 hours from day 0 (iPSC, before treatment with differentiation initiations) to day 15 for every cell line. After each time-course was completed, the RNAs were extracted from frozen cells and sequenced[9]. Hence, each sequenced data corresponds to the average transcription of cells in a well. The procedure was repeated for each of the 19 cell lines, hence each cell line corresponds to an independent differentiation process. To calculate the scEntropy for each cell line, we take the gene expressions at day 0 (the undifferentiated state) as the reference cell for different time points of the same cell line, so that scEntropy gives the relative transcription order at each day with respect to day 0. The scEntropies of the sequenced cells from each cell line at each day are shown in Fig. 3, which also shown an obvious tendency of increasing scEntropies along the differentiation process from day 1 to day 15, and please note that the scEntropies for each cell line at day 0 are zero.

**Figure 3:**
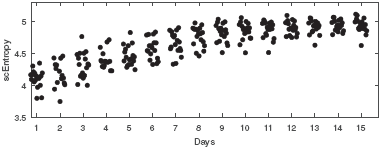
scEntropy dynamics during the differentiation of iPSCs to cardiomyocytes. Each dot gives the scEntropy of the average transcription of cells in a well for each cell line. The dots are scattered randomly in each day for avoiding mixing and overlapping.

### 2.2 scEntropy as a pseudo-time of stem cell differentiation

During cell reprogramming and differentiation, the gene expressions associated with cell pluripotency dynamically change along with cell type transitions. The above examples shown that scEntropy is in general increase with the losing of cell pluripotency, and hence can be used to quantify the process of cell reprogramming and differentiation. Thus, we can consider scEntropy as an intrinsic state variable of the pluripotency of cells, and the increasing of scEntropy as a pseudo-time related to cell reprogramming/differentiation. Here, we analyze the process of kidney organoid differentiation to illuminate how scEntropy may help us to reveal the biological processes of cell type changes.

Consider scEntropy as a pseudo-time, it is straightforward to study how the expressions of each gene varies over the differentiation process. Based on the RNA sequencing data during kidney organoid differentiation[12] (Fig. 1), we calculate the Pearson correlation coefficients of the expressions of each gene with the scEntropy. There are genes that show highly positive correlation with the scEntropy (Fig. 4A). They are potential marker genes of the differentiation process and the gene expression levels change with similar tendency as the transcriptional order (Fig. 4A). The positively correlated genes include the oncogenes KRAS, CTNNB1, RAB14, and PRDX2, the genes SMAD5 and FGFR1 that are involved in the transforming growth factor beta signaling pathway, and the genes that regulate fundamental cellular processes, such as CLIC1 that regulates the cell membrane potential and cell volume, VDAC2 that is involved in the main pathway for metabolite diffusion across the mitochondrial outer membrane, and SRSF6 that is involved in mRNA splicing. Some genes show negatively correlation between gene expression and the scEntropy. Most negative correlated genes encode ribosomal proteins. Scatter plots for gene expression and scEntropies of 20 significant correlated genes are shown in Fig. 4B

**Figure 4:**
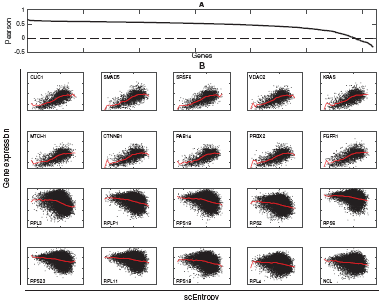
Pearson correlation between gene expression and scEntropy. **A**. Pearson correlation coefficient (PCC) of genes that expressed in at least 80% of cells. There are totally 1230 genes. The genes are ordered according to the PCC value. **B**. Expressions of the 10 positive correlation and 10 negative correlation genes versus the scEntropy. Red lines show the dependence of average expression levels with scEntropy.

In the above discussions, we have considered the scEntropy and gene expression at single cell level. Now, we compare the average scEntropy and gene expression for cells at each day during the differentiation process. Pearson correlation coefficients between the average gene expressions and scEntropies are shown in Fig. 5A. Here, we note that most genes show negative correlations with the scEntropies. This is different from the observation at the single-cell level that most genes show positive correlations. Since the scEntropy can be considered as the pseudo-time of stem cell differentiation, negative correlation implies the decreasing of gene expression along the differentiation process, while positive correlation implies the increasing of gene expression. Fig. 5B shows the relation between Pearson correlation and the fold change of gene expression at day26 with respect to day 0. The Pearson coefficient depends on the fold change in a form of a S-shape function. The density of the fold change also shows that, the average expression level of most genes (83%) at day 26 is lower comparing with that at day 0 (Fig. 5B). These results indicate that scEntropy can be served as a pseudo-time of stem cell differentiation, however the correlation relation between gene expression can be different at single-cell level and at the average level of many cells.

**Figure 5:**
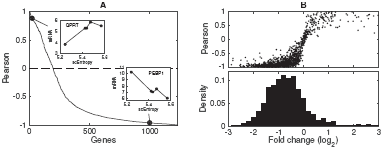
Average scEntropy and gene expression for cells at each day during the differentiation process. **A**. Pearson correction of genes versus average gene expressions. Insets show the relation between average gene expression and scEntropy of the gene QPRT and PEBP1. **B**. Upper panel, Pearson correlation versus fold change of gene expression at day 26 with respect to day 0. Lower panel, density of the fold changes (in log_2_).

### 2.3 Evolution dynamics of the scEntropy distribution

To further investigate the dynamic process of cell date changes, we analyze the distribution of scEntropies of cells during kidney organoid differentiation (Fig. 6A, solid dots). Usually, the evolution dynamics of the distribution can be described by a Fokker-Planck equation of form

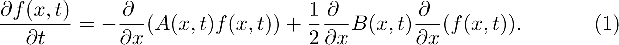

**Figure 6:**
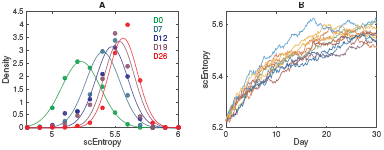
Evolution of the scEntropy distribution during kidney organoid differentiation. **A**. The density of cells at different days during kidney organoid differentiation. Solid dots are obtained from the scRNA-seq data, lines are theoretical distributions (3) based on the Fokker-Planck equation (1). **B**. Simulated scEntropy dynamics based on the stochastic differential equation (4). Here, the parameters are *k* = 0.09 day^−1^, *B* = 0.00108 day^−1^, *x*\*= 5.60, and the initial condition *x*_0_ = 5.23.

Here, *f*(*x, t*) = *P* {*X*(*t*) = *x*} means the probability of a cell to have scEntropy x at time t. To obtain the evolution dynamics of the scEntropy, we need to construct the functions *A*(*x, t*) and *B*(*x, t*) by fitting the solution of equation (1) to the data. From Fig. 1, the average of the scEntropy increase, and the variance decrease with time. Hence, we can try a simple base-line one-attractor model so that[5]

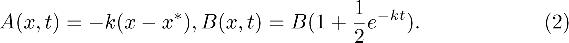

Here *k, x*\*, and *B* are the model parameters.

From the equation (1) and the model (2), we obtain a solution

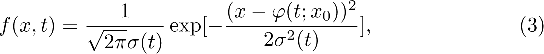

where *x*_0_ is the average scEntropy at *t* = 0, and

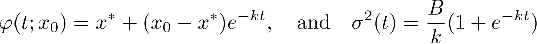

give the average and variance of the scEntropy, respectively. Hence, we obtained the model parameter by fitting ϕ(*t; x_0_*) and *σ*^2^(*t*) to the data in Fig. 1, which gives *x*\* = 5.60; *k* = 0.09 day^−1^, *B* = 0.00108 day^−1^ and *x*_0_ = 5.23. Solid lines in Fig. 6A show the distribution evolution obtained from the Fokker-Planck equations with the obtained parameters.

The Fokker-Planck equation of form (1)-(2) suggests a stochastic differential equation

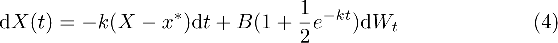

for the evolution dynamics of scEntropy, where *X*(*t*) represents the scEntropy of a cell at time *t*, and *W_t_* is the Weiner process. Given an initial condition *X*(0) = *x*_0_, the solution of (4) gives the random dynamics of the scEntropy of cells with a given initial condition (Fig. 6B). In particular, the time course of the scEntropy density of all cells starting from the initial state *X*(0) = *x*_0_ is obtained by the solutions of (1) with initial condition *f*(*x*, 0) = *δ*(*x* − *x*_0_), which is given as

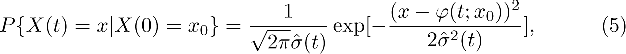

where ϕ(*t*, *x*_0_) is given previously, and

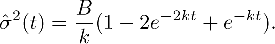

The above arguments suggest that the stochastic differential equation of form (4) provides a way to describe the differentiation process of a single cell through the random scEntropy dynamics. Given the parameters and initial condition, a sample solution of (4) gives a possible trajectory of scEntropy of a single cell during differentiation. Fig. 6B shows the trajectories obtained from (4). Currently, we can only sequence a cell once, and it is impossible to track the evolution of transcriptome for a single-cell. Here, the equation (4) provides a conceptual description of the transcriptional states of a cell during kidney organoid differentiation.

## 3 Discussion

The novel concept of scEntropy has been defined to measure the macroscopic transcription order of individual cells based on scRNA-seq[6]. The concept of scEntropy is valuable in describing the process of early embryo development, as well as the classification between normal and malignant cells in different types of cancers[6]. The current study is the first to introduce scEntropy as a macroscopic quantity of the transcription state of cells to describe the process of iPSCs reprogramming and differentiation. The scEntropy can be a pseudo-time of the differentiation/reprogramming process and show decreasing with the increase of cell pluripotency. The proposed scEntropy is an intrinsic quantify of the cell transcriptional state, and hence the genes that show expression levels correlated with the scEntropy can be essential for the process of cell fate decision. We also consider the scEntropy dynamics of an individual cell during cell fate transition as a stochastic process, which can be modeled through Langevin equations. The conceptual model of Langevin equation provides insights on the dynamics of cell state changes during the differentiation/reprogramming of iPSCs. Based on the stochastic dynamics of the scEntropy, the epigenetic landscape of cell differentiation/reprogramming is described by the corresponding Fokker-Plank equation. Our results manifest the cell-to-cell variability of scEntropy during the differentiation process. Similarly, heterogeneity of gene expression measured by the Shannon entropy also increases in the first few hours in the differentiation process of chicken erythroid progenitors[8] and in the differentiation from pluripotent stem cells to neuronal state[10]. Both entropies from cell- and gene-based show increase during the differentiation process, which suggests the features of variability and stochasticity in stem cell differentiation.

The scEntropy was proposed to measure the intrinsic order of the cellular transcriptome with respect to a predefined reference cell. The definition of scEntropy includes no external parameters, and hence can provide natural information of a cell. It is essential to quantify the changes of intrinsic cellular transcriptome during differentiation/reprogramming for our understanding of the biological process of cell fate decision. iPSCs provide a controllable system to study the process of cell fate changes at individual cell level. Single-cell sequencing techniques enable us to examine the microscopic states of individual cells. However, there are two major limitations when we analyze the single-cell sequencing data: (1) each cell can only be sequenced once and the time course of a cell can not be tracked; (2) the sequencing data are usually very high dimensional, and it is usually difficult to find the consistent low dimensional features to characterize the cells. Hence, for a better understanding of the process of cell type transition, it is important to quantify the cell types through the intrinsic state of a cell. Our study shows that scEntropy can be one of the potent variable for the intrinsic state of cellular transcriptome, and can be used to phenomenally describe the single cell dynamics of epigenetic landscape through the stochastic dynamical models. The current approach can be extended to explore more detailed dynamics of various biological processes of cell type transitions, such as early embryo development, cancer cell plasticity, and cell differentiation.

## Acknowledgments

This work is supported by the National Natural Science Foundation of China (11831015, and 11872084).

## References

[1] Davide Cacchiarelli, Cole Trapnell, Michael J Ziller, Magali Soumillon, Marcella Cesana, Rahul Karnik, Julie Donaghey, Zachary D Smith, Sutheera Ratanasirintrawoot, Xiaolan Zhang, et al. Integrative analyses of human reprogramming reveal dynamic nature of induced pluripotency. Cell, 162(2):412–424, 2015.

[2] GTEx Consortium et al. Genetic effects on gene expression across human tissues. Nature, 550(7675):204, 2017.

[3] Charles Gawad, Winston Koh, and Stephen R Quake. Single-cell genome sequencing: current state of the science. Nat Rev Genet, 17(3):175, 2016.

[4] Lin Guo, Lihui Lin, Xiaoshan Wang, Mingwei Gao, Shangtao Cao, Yuanbang Mai, Fang Wu, Junqi Kuang, He Liu, Jiaqi Yang, Shilong Chu, Hong Song, Dongwei Li, Yujian Liu, Kaixin Wu, Jiadong Liu, Jinyong Wang, Guangjin Pan, Andrew P Hutchins, Jing Liu, and Duanqing Pei. Resolving cell fate decisions during somatic cell reprogramming by single-cell RNA-seq. Mol Cell, 73(4):815–829, 2019.

[5] Qin Li, Anders Wennborg, Erik Aurell, Erez Dekel, Jie-Zhi Zou, Yuting Xu, Sui Huang, and Ingemar Ernberg. Dynamics inside the cancer cell attractor reveal cell heterogeneity, limits of stability, and escape. Proc Natl Acad Sci USA, 113(10):2672–2677, 2016.

[6] Jingxin Liu, You Song, and Jinzhi Lei. Single-cell entropy to quantify the cellular transcriptome from single-cell RNA-seq data. Biophys Rev Lett, doi:10.1142/S179304802500010, 2020.

[7] Dan L Nicolae, Eric Gamazon, Wei Zhang, Shiwei Duan, M Eileen Dolan, and Nancy J Cox. Trait-associated SNPs are more likely to be eQTLs: annotation to enhance discovery from GWAS. PLoS Genet, 6(4):e1000888, 2010.

[8] Angélique Richard, Loïs Boullu, Ulysse Herbach, Arnaud Bonnafoux, Valérie Morin, Elodie Vallin, Anissa Guillemin, Nan Papili Gao, Rudiyanto Gunawan, Jérémie Cosette, Ophélie Arnaud, Jean-Jacques Kupiec, Thibault Espinasse, Sandrine Gonin-Giraud, and Olivier Gandrillon. Single-Cell-Based Analysis Highlights a Surge in Cell-to-Cell Molecular Variability Preceding Irreversible Commitment in a Differentiation Process. PLoS Biol, 14(12):e1002585, 2016.

[9] BJ Strober, Reem Elorbany, K Rhodes, Nirmal Krishnan, Karl Tayeb, Alexis Battle, and Yoav Gilad. Dynamic genetic regulation of gene expression during cellular differentiation. Science, 364(6447):1287–1290, 2019.

[10] Patrick S Stumpf, Rosanna C G Smith, Michael Lenz, Andreas Schuppert, Franz-Josef Müller, Ann Babtie, Thalia E Chan, Michael P H Stumpf, Colin P Please, Sam D Howison, Fumio Arai, and Ben D MacArthur. Stem cell differentiation as a non-Markov stochastic process. Cell Syst, 5(3):268–282.e7, 2017.

[11] Kazutoshi Takahashi, Koji Tanabe, Mari Ohnuki, Megumi Narita, Tomoko Ichisaka, Kiichiro Tomoda, and Shinya Yamanaka. Induction of pluripotent stem cells from adult human fibroblasts by defined factors. Cell, 131(5):861–872, 2007.

[12] Haojia Wu, Kohei Uchimura, Erinn L Donnelly, Yuhei Kirita, Samantha A Morris, and Benjamin D Humphreys. Comparative analysis and refinement of human PSC-derived kidney organoid differentiation with single-cell transcriptomics. Cell Stem Cell, 23(6):869–881, 2018.

[13] Zhihong Zhu, Futao Zhang, Han Hu, Andrew Bakshi, Matthew R Robinson, Joseph E Powell, Grant W Montgomery, Michael E Goddard, Naomi R Wray, Peter M Visscher, et al. Integration of summary data from GWAS and eQTL studies predicts complex trait gene targets. Nat Genet, 48(5):481, 2016.

